# CXCR6 by increasing retention of memory CD8 T cells in the ovarian tumor microenvironment promotes immunosurveillance and control of ovarian cancer

**DOI:** 10.1101/2020.12.02.401729

**Authors:** Ravikumar Muthuswamy, AJ Robert McGray, Sebastiano Battaglia, Wenjun He, Anthony Miliotto, Cheryl Eppolito, Brian Lichty, Protul Shrikant, Kunle Odunsi

**Author notes:** **Corresponding Authors:** Ravikumar Muthuswamy, Roswell Park Comprehensive Cancer Center, Elm and Carlton Streets, Buffalo, NY14263, Kunle Odunsi, MD, PhD, <u>Current Corresponding address</u>: University of Chicago Medicine Comprehensive Cancer Center, 5841 S Maryland Ave, Chicago, IL 60637.

## Abstract

**Purpose:** Resident memory CD8 T cells owing to their ability to reside and persist in peripheral tissues, impart adaptive sentinel activity and amplify local immune response, have beneficial implications for tumor surveillance and control. The current study aims to clarify the less known chemotactic mechanisms that govern the localization, retention, and residency of memory CD8 T cells in the ovarian tumor microenvironment.

**Experimental Design:** RNA/FACS based profiling of chemokine receptor expression in CD8^+^ resident memory T cells in human ovarian cancer and analyze their association with survival. Analyze chemokine receptor role in anti-tumor response and control by resident memory T cells using prophylactic mice models of ovarian cancer, treated with adoptive transfer of OT1 T cells and vaccination with maraba virus-OVA to target Ovalbumin expressing tumor.

**Results:** Chemokine receptor profiling of CD8^+^CD103^+^ resident memory TILs in ovarian cancer patients revealed high expression of CXCR6. Analysis of the TCGA ovarian cancer database revealed CXCR6 to be associated with CD103 and increased patient survival. Functional studies in mouse models of ovarian cancer revealed that CXCR6 is a marker of resident, but not circulatory tumor-specific memory CD8 T cells. Knockout of CXCR6 in tumor-specific CD8 T cells showed reduced retention in tumor tissues leading to diminished resident memory responses and poor control of ovarian cancer

**Conclusions:** CXCR6 by promoting increased retention in tumor tissues serves a critical role in resident memory T cell-mediated immunosurveillance and control of ovarian cancer. Future studies warrant exploiting CXCR6 to promote resident memory response in cancers.

## Introduction

In ovarian cancer patients, the density and location of CD8^+^ tumor-infiltrating lymphocytes (TILs) are critical for determining progression-free and overall survival (1,2). However, recent studies indicate that only ~ 10% of ovarian tumor-infiltrating lymphocytes (TILs) are specific for shared antigens or mutated neoantigens (3). Moreover, the phenotype of tumor-specific T cells, rather than simply the sheer number of T cells is a crucial parameter in determining effective cancer immunity (3–5). In this regard, eliciting T cell responses with durable memory attributes in tumors can lead to better clinical outcomes in cancer patients (6,7). Tissue-resident memory (Trm) cells are a unique subset of memory T cells that lose their recirculatory potential and take up residency in peripheral tissues in areas of previous antigen encounter (8–10). This contrasts with central memory T cells (TCM) that predominantly circulate within secondary lymphoid organs (SLO) and effector memory T cells (TEM), which recirculate mostly in peripheral tissues. Trm cells are considered as an amalgamation of effector memory TEM and TCM as they share features with both subsets, having effector molecules similar to TEM and renewal and longevity potential similar to TCM (11). Owing to their residency in peripheral tissues, they can act as adaptive sentinels capable of inducing rapid and robust antigen recall responses, by eliciting systemic effector functions, without the requirement for priming in the lymph node (12–14). They are metabolically adept to persist and survive longer in peripheral tissues (15). Further, studies in mice and humans indicate their presence in tumors predicts enhanced tumor control and progression-free survival (16–18).

Based on these attributes, Trm cells represent an important T cell subset capable of generating robust anti-tumor immunity, and therapies like adoptive cell transfer and immune checkpoint blockade will greatly benefit from improving their accumulation in tumors (19,20). However, little is known about the factors that govern Trm localization and retention in tumor tissues, one of the critical steps in the generation of Trm responses. Thus, identifying the factors and mechanisms that drive Trm localization and retention has profound therapeutic implications for promoting Trm response in tumors, as there are no strategies currently available to promote Trm response in cancer patients.

Chemokines and chemokine receptors are implicated in the mobilization and localization of immune cells to peripheral tissues and the generation of memory response (21–23). While a role for CXCR6 in CD8+ T cell resident memory responses has been reported in infectious disease models (24–26), there are no studies that directly address if specific chemokine receptors facilitate resident memory response by driving localization or retention of CD8^+^ T cells in cancerous tissues. Our current study addresses this gap and identified CXCR6 as the predominant chemokine receptor expressed by Trm CD8^+^ T cells in human ovarian cancer. Using murine models of ovarian cancer, we demonstrate that CXCR6 marks tumor-specific resident memory T cells, and their presence in tumors is associated with increased survival. The deletion of CXCR6 in tumor-specific CD8^+^ T cells resulted in reduced retention in tumor tissues and increased CD8^+^ T cell recirculation to the spleen, culminating in diminished resident memory response and reduced control of ovarian tumors. These findings indicate that CXCR6 is required for efficient generation of tumor-specific resident memory T cell responses and could be therapeutically exploited for control of ovarian cancer.

## Materials and Methods

### Human studies

Tissue samples were collected from patients undergoing primary tumor debulking surgery for ovarian cancer under a protocol approved by the institutional review board (protocol # I215512). For correlative analysis of immune markers and patient survival, a publicly available ovarian cancer TCGA Firehose legacy 2020 mRNA-seq database (n=307) was used. Out of this, only high-grade (Stage IIIA to IV, N=280) patients were used for further analysis (Suppl. Fig 1A). Correlation and survival analysis were done in either whole set (N=280) or in a subset of patients stratified for high CD8 levels (top 75 percentile N=210 was used, while the bottom 25 percentile, N=70 was not included). For survival analysis with CXCR6, a comparison between the top and bottom 25 percentile was done either in whole or patients stratified for high CD8.

### Mice studies

Wild type (Wt.) C57BL/6 and CXCR6KO were purchased from The Jackson Laboratory (Bar Harbor, ME). OT1 PL RAGKO (Mice that expresses TCR for OVA_257-264_ peptide and has no endogenous CD4^+^, CD8^+^, T, and B cells due to knockout in RAG gene) was a kind gift from Dr. Shrikant. Wt. C57BL/6 mice were expanded by breeding. OT1 PL RAGKO mice were crossed with CXCR6KO to generate OT1 mice that were deficient for CXCR6. All the above crossing and expansion was done under breeding protocol 1145M and experiments involving the above mice were performed as per the 1371M experimental protocol. All animals were maintained in the Laboratory Animal Shared Resource under pathogen-free conditions in the Laboratory Animal Resource (LAR) facility located in Roswell Park Comprehensive Cancer Center (RPCCC). All animal experiments were carried out according to protocol guidelines reviewed and approved by the Institute Animal Care and Use Committee (IACUC) of Roswell Park Comprehensive Cancer Center (Buffalo, NY).

### Adoptive Cell transfer

OT1 CD8^+^ T cells from Wild type or the knockout mice mentioned above were purified from spleens using EasySep™ Mouse CD8+ T Cell Isolation Kit (#19853, STEMCELL Technologies, Vancouver, CA). Isolated CD90.1^+^ OT1 CD8^+^ T cells were injected into CD90.2^+^ C57BL6 recipient mice at a concentration of 10^6^ cells/100ul PBS through the retro-orbital route.

### Vaccination

Attenuated strain MG1 of Maraba virus (27,28) expressing full-length ovalbumin (OVA)was manufactured at McMaster University, their titer determined, shipped on dry ice to RPCCC, and stored at −□80□°C before use. They were injected at 10^7^ PFU intraperitoneally for use as vaccine/adjuvant to activate OT1 T cells, that were adoptively transferred one day before to C57BL6 mice.

### Tumor model

IE9-mp1, a derivative of widely used ID8 mouse ovarian surface epithelial cell line (29) obtained by one passage in mice and genetically engineered to express Ovalbumin(30), was the tumor cell line used for *in vivo* mice study. Cell line was cultured in RPMI-1640 (10-040-CM) + 10% fetal bovine serum (35-010-CV) in presence of 1x penicillin/streptomycin (100 IU /100 ug mL^−1^, 30-002-CI) at 37°C, 5% CO2 conditions. All reagents used for cell line culture are from CORNING, Corning, NY. BL6 mice were injected intraperitoneally (I.P.) with 1× 10^7^ cells of IE9-mp1. All the control and treated mice were euthanized when their abdominal circumference is ≈10 cm, which was considered as the experimental endpoint.

### Real-time qPCR analysis

RNA from tumor and cells were isolated by RNeasy kit (#74106, Qiagen,). cDNA conversion of isolated RNA was done by using isCript™ cDNA synthesis kit (#170889, BIO-RAD). cDNA was quantified by using predesigned KiCqStart SYBR^®^ Green primers (# KSPQ12012, Sigma Aldrich, sequences of the primers provided in supplementary table-1,) specific for the various human chemokine receptor genes and IQ™ SYBR green reagent (# 170882, BIO-RAD) on CFX96 real-time PCR detection system (BIO-RAD). QPCR data were analyzed on CFX manager 3.1 (BIO-RAD).

### Flow cytometry of human and mice samples

Human tumors were minced into small pieces, suspended in RPMI media, homogenized into single-cell suspension using Miltenyi gentleMACs dissociater. The single-cell suspension was passed through a 100micron cell strainer, washed, spun on ficoll to remove debris and dead cells. Tumor-infiltrating lymphocytes (TILs) processed from above were further processed for downward analysis. For flow-based sorting of human Trm and non-Trm cells, above processed TILS, were stained for live/dead dye (Zombie Aqua fixable viability kit, #423102 BioLegend) in PBS, followed by staining with BioLegend antibodies specific for CD3(Pe-Cy7, HIT3a, #300316), CD8(BV421, RPA-T8, #300106), CD103 (Alexa Flour 488, Ber-ACT8, #350208) and sorted on BD FACSAria™ II sorter into CD8^+^ CD103^−^ (Non-Trm) and CD8^+^ CD103^+^ (Trm) cells and further processed for RNA analysis. Mice tumors were similarly processed, except for the ficol step, and were straightaway stained with flow antibodies. Human tumor samples were stained with live/dead dye in PBS, followed by staining with BioLegend antibodies specific for CD3, CD8, CD103, CCR5 (APC-cy7, J418F1, #359110,) and with BD biosciences antibodies specific for CXCR3 (PE, IC6, #560928), CXCR4(APC,12G5, #560936) CXCR6(BV786, 13B 1E5, #743602) and in FACS buffer. Mice tissue samples were stained with live/dead dye, followed by antibodies specific for CD8(FITC, 53-6.7, #553031), CD90.1 (PE, OX7, 554898), CD103 (A700, M290, 565529) from BD biosciences, CXCR6 (BV421, SAO51D1, #151109), CD44 (APC-Cy7, IM7, #103208), CD62L(BV786, MEL-14, 104440), CD69(PE-Cy7, H1.2F3, #104512), CXCR3 (BV650, CXCR3-173, #126531) from BioLegend. Stained samples were acquired on the BD LSRII flow cytometry system using BD FACSDiva software. Post-run analyses were done using the FCS Express 7 research edition software (De Novo Software, Pasadena, CA).

### Confocal microscopy

Human or mouse tissue sections were marked on the edge with ImmEdge pen (H-4000, Vector Labs) and fixed in 4% PFA, 15 minutes. Human and mouse tissues were blocked with 5% human or mouse serum respectively for 1 hour at room temperature (RT). This was followed by Incubation with fluorophore-conjugated (1:100 dilution) or non-conjugated primary antibodies at (1/500 dilution) for 3 hours at RT. Washed 4 times in 1x PBS. To visualize tumor antigens targeted with non-conjugated primary antibodies, fluorophore-conjugated secondary antibodies(1:1000 dilution, Cell signaling technology) were added along with nuclear dye (Hoechst 33422, # H1399, Thermo Fisher Scientific) for 30 minutes at RT. Washed 4 times in 1x PBS and sections are mounted with coverslips in prolong gold mounting medium. Tissues are analyzed in Leica SP8 DMRE spectral confocal microscope and Image J software was used for post-image analysis. The full details of antibodies and secondary antibodies used for confocal staining of human and mouse tissues are given in supplementary table 2.

### Cytotoxic T lymphocyte induced activated Caspase3/7 assay

5 × 10^5^ Wt. OT1 and CXCR6 OT1 were incubated by itself or with 5 × 10^4^ of OVA expressing IE9-mp1 target cell line or OVA non expressing ID8 cell in the presence of 4uM CellEvent™ Caspase-3/7 Green Detection Reagent (#C10423, ThermoFischer Scientific) in triplicates for 24 hours. Caspase-3/7 fluorescence is measured on Biotek Synergy HT microplate plate reader at emission 528nM. Images of the wells with each condition were taken on ZOE™ fluorescent cell imager (BIO-RAD).

### Statistical analysis

All statistical analyses involved in the study were performed using GraphPad Prism 8 software. The type of statistic used is described in the figure legends respectively. For all analyses, P<0.05 was considered statistically significant. *, **, ***, and **** indicates P<0.05, P<0.01, P<0.001 and P<0.0001 respectively. All Data were expressed as Mean ± SEM except in some cases, where box and whisker plots were used. In the box and whisker plots, the horizontal line represents the median and the whiskers connect the highest and lowest observations. Log-rank (Mantel-Cox) test was used to calculate the mice survival.

## Results

### Chemokine receptor CXCR6 is highly expressed on human tumor resident memory CD8+ T cells

To characterize the potential chemokine/chemokine receptor axes involved in the localization of tissue-resident memory CD8^+^ T cells in the ovarian cancer tumor microenvironment (TME), we profiled the expression of the known chemokine receptors in Trm (CD3^+^CD8^+^CD103^+^) vs. non-Trm (CD3^+^CD8^+^CD103^−^) cells obtained from ovarian cancer patient tumors using RT-PCR and flow cytometry. We used CD103 for identifying Trm cells, as it is considered a canonical marker for tissue-resident memory cells and predicts favorable outcomes in ovarian cancer patients (31,32). While mRNA expression of most chemokine receptors demonstrated no clear differences between Trm and non-Trm, Trm cells consistently showed significant upregulation of CXCR6 mRNA compared to non-Trm Cells (Fig.1A). CCR7 mRNA showed a reverse trend with more expression in non-Trm cells (Fig.1A). Flow cytometry analysis further confirmed the high expression of CXCR6 protein in CD8^+^CD103^+^ Trm cells (Fig. 1B), in agreement with previous reports (33). Although CXCR3 was high in CD103^+^ Trm cells no significant difference was noted in its expression when compared to non-resident CD8^+^ T cells. Analysis with confocal microscopy further substantiated the co-expression of CXCR6 and CD103 on CD8^+^ TILs (Fig.1C).

**Figure 1.**
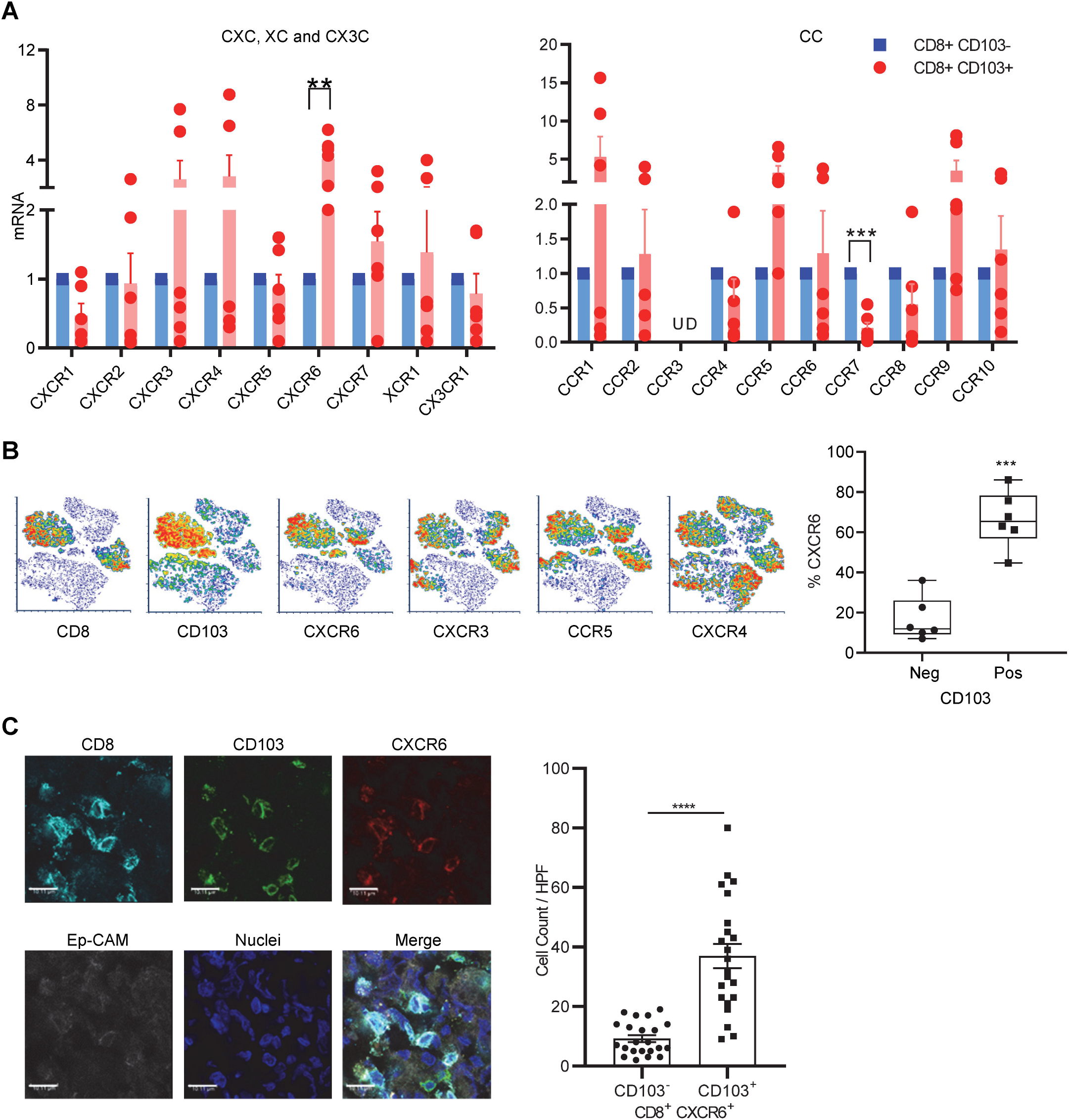
Chemokine receptor CXCR6 is a valid marker of human tumor resident CD8 T cells. (A) CD8^+^TILs from ovarian cancer patients (N=6) were flow-sorted into CD103^+^ (Trm) and CD103^−^(non-Trm) subsets and real-time RT-QPCR was used to profile the expression of CXC, XC & CX3C (Left), and CC (Right) chemokine receptors. mRNA levels were normalized to HPRT1 and chemokine mRNA levels in the CD103^+^ subset were expressed as fold change over CD103^−^ subset. Data presented as Mean±SEM and U.D. denotes undetectable. (B) CD8^+^ TILs in human ovarian cancer were flow stained for CD103 and chemokine receptors and presented in the left panel as TSNE plot from one representative patient and in the right panel as Box and whisker plot of % CXCR6 in CD103^−^ and CD103^+^ CD8 TILs from 6 ovarian cancer patients. (C) Left, images from confocal staining for CD8^+^CD103^+^CXCR6^+^ Trm cells in ovarian cancer (Scale bars are at 10 μM) and right, Bar graphs represent 22 high power field CD103 pos and neg cell counts in CD8^+^CXCR6^+^ TILs from 5 patient tumors. A paired 2 tailed T-test was used in (A), (B), and (C), and **, *** and **** indicate P<0.01, P<0.001, and P<0.0001, respectively.

### TCGA analysis reveals a positive correlation of CXCR6 with CD103 and survival of ovarian cancer patients

Analysis of the whole database of high-grade serous ovarian cancer patients (N=280) or patients stratified for high CD8 (N=210) showed a significant correlation between CD103 and CXCR6 mRNA markers (Fig. 2A). Details of TCGA database stratification are provided in supplementary figure-1A-C. Furthermore, survival analysis using TCGA data revealed that the CXCR6 marker to be positively associated with increased survival in CD8 high but not in the whole dataset (Fig 2B).

**Figure–2.**
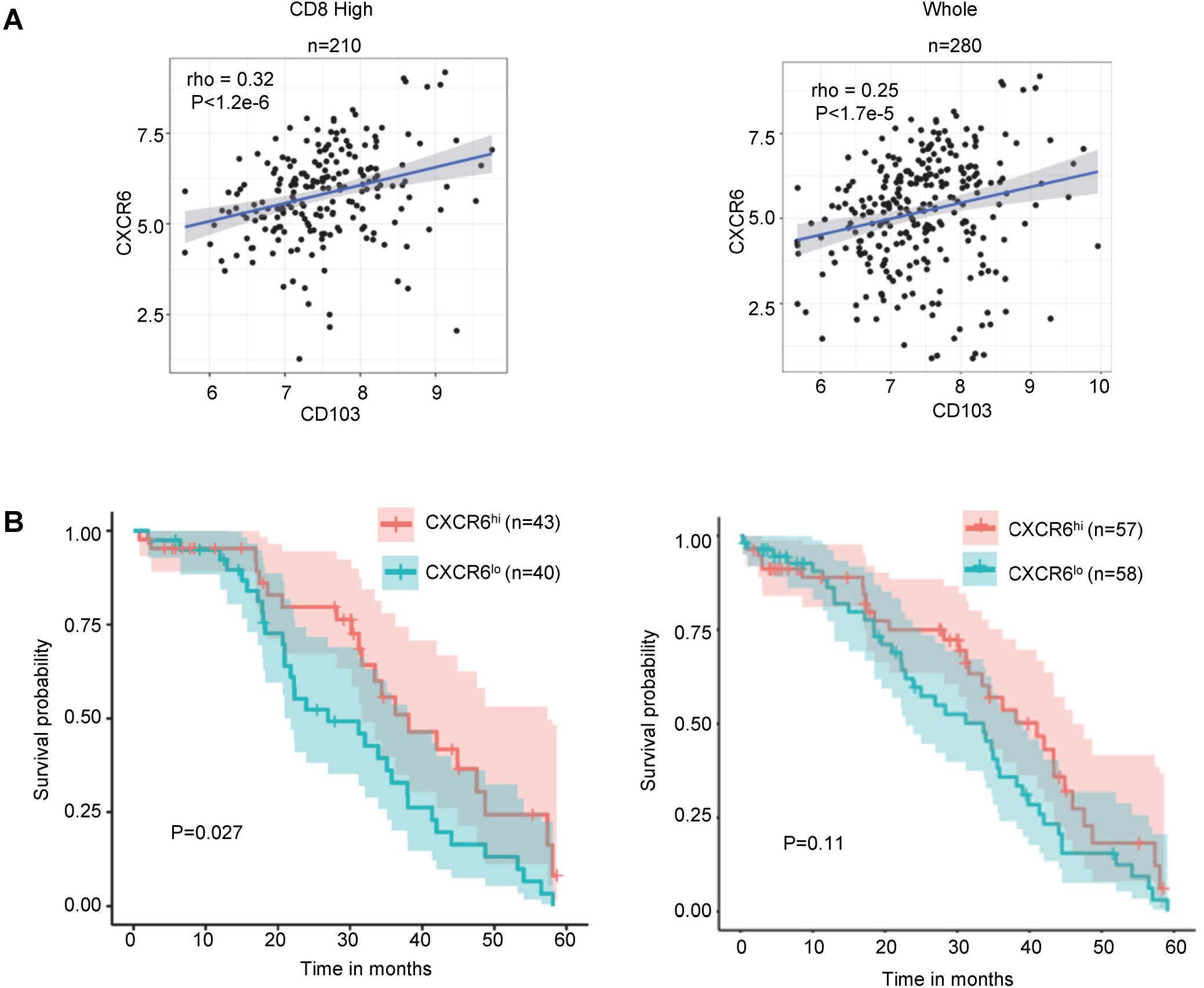
CXCR6 shows a high correlation with CD103 and protection against ovarian cancer based on Ovarian TCGA database analysis. (A) Spearman’s rank correlation between CXCR6 and CD103 markers was done in the CD8 high (left, N=210) or the whole set (N=280) of high-grade TCGA ovarian patient database. Analysis of association of CXCR6 (B) markers with survival in the CD8 high (left) or the whole set (right) of TCGA database of high-grade ovarian patients.

### Mice treated with adoptive transfer of OT1 and vaccination with Mrb-OVA treated mice display higher survival

To directly test whether CXCR6 plays a functional role in the localization and efficacy of tumor resident memory responses, we developed a prophylactic vaccination model using a mouse intraperitoneal ovarian cancer model as previously described (34,35). We used the prophylactic model instead of the therapeutic setting for the following reasons: (i) the continued presence of tumors may promote more of the effector rather than memory responses, (ii) response to the tumor may not necessarily derive from memory CD8 T cells and (iii). the OVA expressing IE9-mp1 tumor model is highly aggressive, and the cure rate is low (35), which makes priming and challenge with the tumor difficult. This model (Fig.3A) utilizes adoptive cell transfer (ACT) of naïve OT1 CD8^+^ T cells into B6 recipient, followed by priming with OVA expressing maraba virus (Mrb-OVA) vaccination to activate OT1 cells. We chose OT1 T cells for adoptive transfer to allow for careful monitoring and tracking of T cell responses as shown in previous studies (13,36). OVA expressing Maraba virus was utilized as the vaccine as it was shown to induce robust anti-tumor CD8^+^ T cell responses in our previous study (35). Tumor challenge was done 31-30 days after adoptive transfer of OT1 and vaccination with Mrb-OVA. This is to give enough time for the vaccine-induced effector response to subside and allow memory generation. This increases the probability that anti-tumor immunity following tumor challenge would come from vaccine-induced memory cells. In this model, the OT1 + Mrb-OVA combination provides better tumor control than individual treatments with either OT1 or Mrb-OVA, (Suppl. Fig. 2A-C). As shown in Fig. 3B, the combination of ACT and vaccination with Mrb-OVA improved tumor protection compared to no treatment as indicated by a median survival of 70 vs. 33 days.

**Figure–3.**
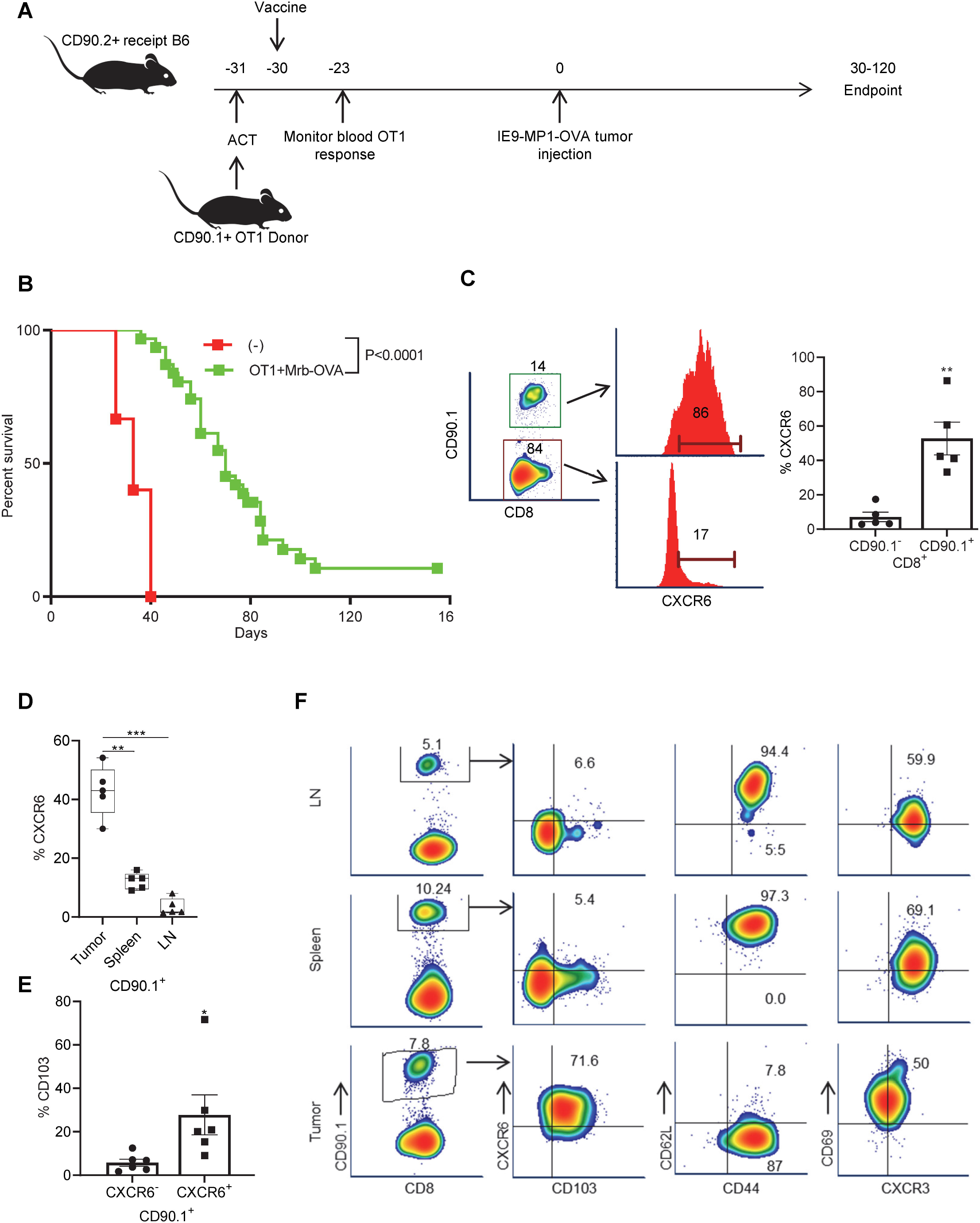
OT1+Mrb-OVA treated mice display higher survival and CXCR6 defines tumor-specific resident memory CD8 T cells in Mice models of ovarian cancer. (A) The experimental schema for adoptive cell transfer of OT1 cells in conjunction with Mrb-OVA vaccination is used to drive immune response against OVA expressing IE9-mp1 tumor cell line. (B) Comparison of survival between mice treated with PBS and with OT1+Mrb-OVA combination. Mice are N=31 for OT1+Mrb-OVA and N=15 for control. Log-rank (Mantel-Cox) test was used to calculate the mice survival and median survival is 70 for OT1+Mrb-OVA and 33 for control groups. (C). CXCR6 expression in non-tumor (CD90.1^−^) and tumor-specific (CD90.1^+^) CD8^+^ T cells in tumors of mice treated with adoptive cell transfer of CD90.1^+^ OT1 and Maraba-OVA vaccination (N=5). (D) Comparison of CXCR6 expression in CD90.1^+^ tumor-specific CD8 T cells present in tumor, spleen, and lymph nodes tissues (N=5) and shown as box and whisker plot. (E) Bar graphs of %CD103 in CXCR6^−^ and CXCR6^+^ subsets of OT1 CD8^+^ T cells in tumors tissues of OT1+Mrb-OVA treated mice (N=6). (F) FACS density plots of CD103 and other memory markers expression in CD90.1 gated CD8 T cells in lymph nodes, spleen, and tumors of one representative OT1+Mrb-OVA treated mice. A two-tailed paired T-test was used to analyze data in (C), (D), and (E), and *, **, and *** indicates P<0.05, P<0.01, and P<0.001, respectively. Data in bar graphs (C & E) are Mean+SEM.

### CXCR6 is highly expressed in antigen-specific CD8^+^ T cells that reside in the tumor, but not those in circulation

Analysis of endpoint tumors from mice that were treated with OT1+Mrb-OVA showed that CXCR6 expression was seen predominantly in tumor-specific CD90.1^+^ OT1 (Fig.3C), but less frequently in CD90.1^−^ CD8^+^ T cells. Similarly, analysis of endpoint tumors in Mrb-OVA alone treated mice (Suppl. Fig. 2), revealed that the endogenous tumor-specific CD8^+^ T cells (marked by OVA tetramer staining) showed high CXCR6 expression compared to non-specific (OVA tetramer negative) T cells (Suppl. Fig.3A). When CXCR6 expression was compared between OT1^+^ CD8^+^ T cells in tumor, spleen, and lymph node tissues of OT1+Mrb-OVA treated mice that had reached endpoint, only tumor-infiltrating OT1^+^ T cells showed high CXCR6 expression (Figs.3D and 3F). Analysis of CD103 in CD90.1^+^ TILs, revealed high expression by CXCR6^+^ cells, rather than in the CXCR6^−^ subset (Figs.3E and 3F).

CD44 and CD62L marker-based stratification show that CD90.1^+^ TILs are predominantly of CD44^+^ CD62L^−^ effector memory phenotype, whereas those in lymph nodes and spleen were more of central memory type (Fig. 3F). The observation that Trm cells resembled more of the effector/effector memory than the central memory phenotype, is in agreement with previous observations (17,37,38). Sequential comparison of OT1 CD8^+^ T cells prior to transfer (Top), 7 days post transfer in recipient blood (Middle) and in endpoint tumors (lower) revealed again that CXCR6 is acquired by T cells only in tumors and marks CD44^+^CD62L^−^ CD103^+^ tumor resident memory cells, but not circulatory cells (Suppl. Fig.3B).

Analysis of additional Trm markers, namely CD69 and CXCR3 revealed that they were found in both circulatory and resident memory CD90.1^+^ CD8^+^ T cells (Fig.3F and Suppl. Fig.3B). This discrepancy between the current study and other studies concerning CD69 as a Trm marker can be attributed to the different tissue microenvironment as suggested by Walsh et al.,(39). Uniform expression of CXCR3 on both resident memory and circulatory memory cells in murine studies, further supported our observations in human Trm. Unvarying expression of CXCR3 on both the subsets and selective late expression of CXCR6 on resident memory cells suggests that while CXCR3 may direct default peripheral migration of both memory subsets to the tumor, CXCR6 selectively promotes tissue residency of memory cells. The above observations strongly attest to the validity of CXCR6 as a Trm marker with functional implications in resident memory response to ovarian tumors.

### Cells negative for CD45 and EpCAM contribute more to CXCL16 expression than hematopoietic or epithelial cells to help Trm cells localize within the ovarian tumor microenvironment

As CXCL16 is the main identified ligand for CXCR6 (40), we wanted to identify the cells that produce the CXCL16 in the mice ovarian tumor microenvironment. We stained tissues from endpoint tumors from OT1+Mrb-OVA treated mice with antibodies for mouse CXCL16 and EpCAM (Tumor Cell marker) and CD45 (immune cells). Confocal microscopy analysis (Fig. 4A) revealed multiple sources of CXCL16, as revealed by CXCL16 (Green) staining in EpCAM^+^ (grey), CD45^+^ (red), and in the cells that were negative for EpCAM and CD45. However quantitative analysis revealed that the later EpCAM^−^CD45^−^ cells were the dominant producers of CXCL16, as they contributed to 72% of CXCL16 expression compared to EpCAM^+^ and CD45^+^ cells that contributed 13 and 14% respectively (Fig. 4A). Additional analysis revealed that F4/80+ cells were the major producers of CXCL16 among CD45+ cells, as they contributed about 67% to CXCL16 expression (Fig.4B). To further substantiate the CXCL16 role in Trm localization, We stained for F4/80, CD103, and CXCL16 markers in endpoint tumor tissues and analyzed them using confocal immunofluorescence microscopy. The analysis revealed that most CD103^+^ T cells were found proximal to CXCL16 positive cells (Fig.4C), with most of them within 20uM distance. The above observations suggest that multiple sources of cells express CXCL16 in the ovarian cancer microenvironment to support the local accumulation of CXCR6^+^ Trm cells.

**Figure–4.**
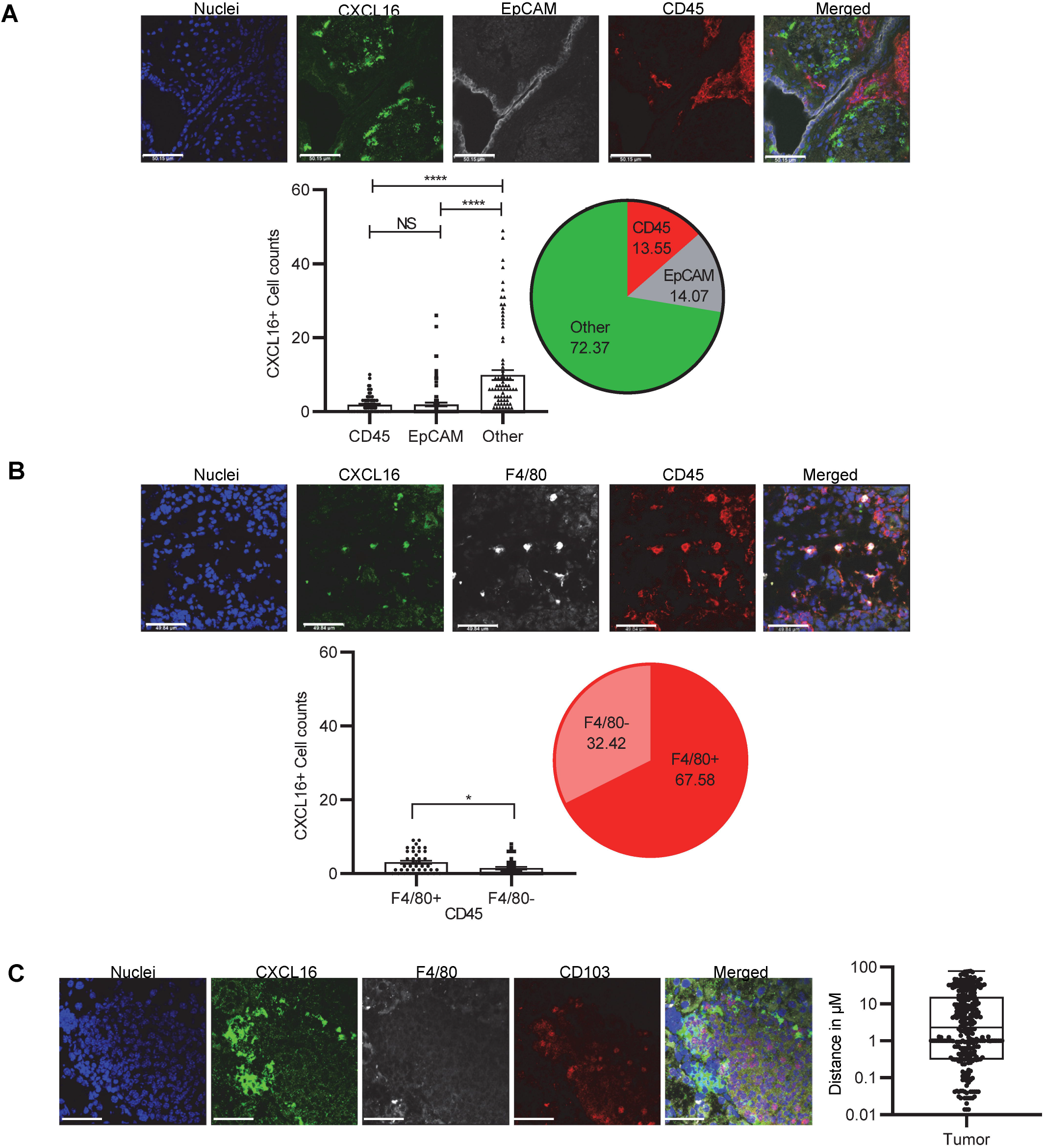
Cells negative for CD45 and EpCAM contribute more to CXCL16 expression to help localize Trm cells in mice ovarian tumor microenvironment. Endpoint tumors from OT1+Mrb-OVA treated mice were stained for nuclei (blue), CXCL16 (Green), along with (A) CD45 (Red) and EpCAM (Grey) or with (B) CD45 (Red) and F4/80 (Grey) or with (C) F4/80 (grey) and CD103 (Red). Bar graphs for (A) represent the total cumulative of 84 high power field (63x magnification) cell counts from 16 tumors, whereas for (B), it is the total cumulative of 40 high power field (63x magnification) cell counts from 15 tumors. The pie diagram represents % contributions from each cell type calculated based on cell counts. in (C) represents 296 distance measurements between cells positive for CD103 and CXCL16 from 5 mice tumors. Scale bars are at 50μM for (A), (B), and (C).

### Knockout of CXCR6 in tumor-specific CD8^+^ T cells enhances circulatory but reduces resident memory response in tumors, leading to diminished protection against ovarian cancer

To determine if CXCR6 plays a critical functional role in Trm generation and tumor immunity, CXCR6KO OT1 mice were generated by crossing OT1 PL RAGKO mice with CXCR6KO mice. *In vitro* testing confirmed no clear phenotypic or functional differences between Wt. and CXCR6KO OT1 T cells, as no differences were noted in marker expression (Suppl. Fig.4A) or their lytic capacity against IE9-mp1 target cells (Suppl. Fig.4B). ACT of Wild type (Wt.) or CXCR6KO (KO) CD90.1^+^ OT1 cells into B6 recipient mice (CD90.2^+^) was performed on Day −31, followed by vaccination with Mrb-OVA at Day −30 and tumor implantation on day 0 (Fig.5A). Blood analysis on Day −23 (7 days after ACT+ vaccination) revealed the B6 recipients that received the adoptive transfer of KO T cells showed significantly higher circulatory T cell response than those that received Wt. T cells (Fig.5B). This was backed by similar findings in the spleen of B6 recipient mice that received KO T cells, as they again showed a higher percentage of CD90.1^+^ T cells compared to Wt. recipients (Fig.5C and Suppl. Fig.5). In contrast quantification of TILs in endpoint tumors, revealed that B6 recipients of CD8 T cells from KO mice had few or no detectable CD90.1^+^ TILs compared to that received from Wt. mice, where CD90.1^+^ TILs were consistently detected (Fig.5D). This was also true for CD103^+^ TILs, as they were also at low frequency or undetectable in recipient mice that received knockout compared to Wt.OT1 (Fig.5E). This difference in transferred T cell persistence in tumors was reflected by differences in tumor progression (Fig.5F) and survival (Fig.5G), as B6 recipients of Wt. CD8 T cells controlled tumors significantly better than KO T cell recipients. Together, the observations confirm that loss of CXCR6 reduces resident memory responses and the associated protection against ovarian cancer.

**Figure 5.**
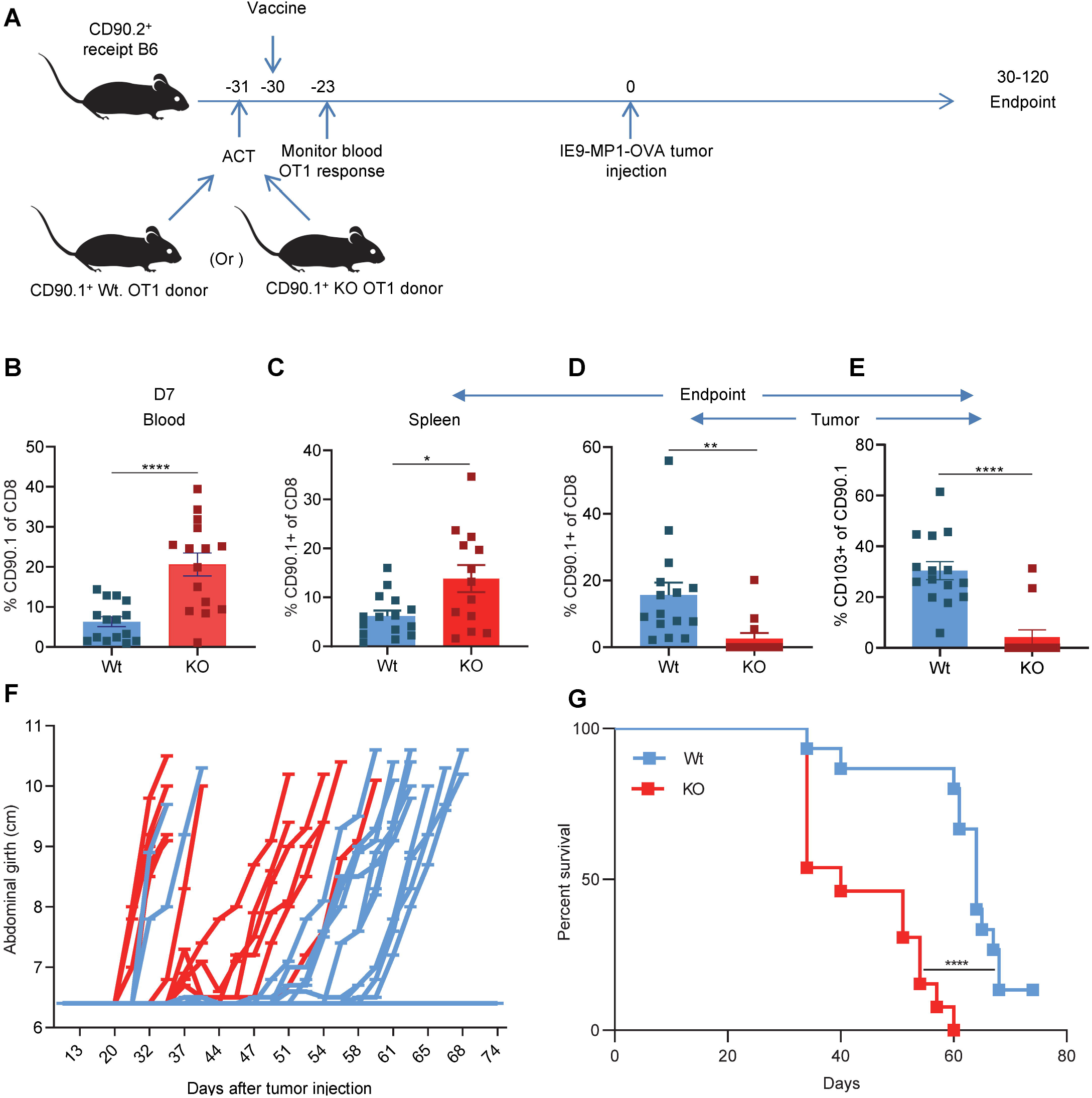
CXCR6KO enhances T cell response in blood, but conversely lowers it in tumor tissues and weakens ovarian tumor control in recipient mice. **(A)** The experimental schema for adoptive cell transfer of Wild (Wt.) or CXCR6KO (KO) OT1 cells in conjunction with Mrb-OVA vaccination in B6 recipient mice. **(B)** % CD90.1^+^ T cells in day 7 post-vaccination blood of B6 recipient mice that received Wt. or KO OT1 cells, N=15 for each. Comparison of % CD90.1^+^ T cells in the spleen **(C),** tumor **(D),** CD103^+^ T cells in the tumor **(E)** at the endpoint, tumor progression **(F),** and survival **(G)** between the 2 groups of B6 mice that received adoptive cell transfer of either Wt. (N=15) or KO (N=13) OT1 cells + Mrb-OVA vaccination. Non paired two-tailed T-test was used to analyze data in **(B)**, **(C), (D),** and **(E),** and data are presented as Mean+SEM. Log-rank (Mantel-Cox) test was used to calculate the survival between mice belonging to Wt. and KO groups in **(G)** with the median survival of 64 and 40, respectively. *, **, *** and **** indicates P<0.05, P<0.01, P<0.001 and P<0.0001, respectively.

## Discussion

The underpinning chemotactic mechanisms guiding Trm localization and retention in the tumor microenvironment (TME) and preventing their emigration have not been clearly defined. Several mechanisms including induction by specific subsets of DC, expression of specific transcriptional regulators HOBIT, BLIMP, Runx3, acquisition of integrins, exposure to homeostatic cytokines and inflammatory signals, and improved metabolic fitness have all been implicated in Trm generation and persistence (15,37,41–44). However, most studies have utilized infectious models and the relevance of these factors and associated mechanisms for the generation of Trm responses in cancer is less understood. Given the beneficial role of Trm in tumor control and the increasing need to develop strategies to promote Trm response in tumors, understanding how Trm response is generated and maintained in tumors is warranted. Specifically, identifying the chemotactic mechanisms that drive Trm response to tumors, will greatly help in the design of immunotherapeutic strategies to promote Trm in patients’ tumors, potentially leading to improved therapeutic efficacy and clinical outcome. Our studies in both humans and mice strongly indicate that CXCR6 is highly expressed on resident memory T cells, plays a critical role in their localization within tumor tissues, and impart increased protection against ovarian cancer. While a role for CXCR6 in CD8^+^ T cell resident memory responses using infectious disease models (24–26) has been reported, to date, there are no reports on the functional role of CXCR6 in mediating Trm responses to tumors. This is critical as most mechanisms underpinning immunity against infectious agents versus tumor challenge are distinct. To our knowledge, this is the first study to demonstrate the indispensable requirement for CXCR6 in tumor resident memory responses and the role in mediating ovarian cancer immunity. Several lines of evidence from our studies support the requirement for CXCR6 in mediating Trm responses, as discussed below.

Our chemokine receptor profiling studies of human ovarian cancer TILs using qPCR, FACS, and confocal analysis all revealed high expression of CXCR6 on CD8^+^ CD103^+^ resident memory CD8 T cells. Analysis of the TCGA database of ovarian cancer patients further corroborated this strong association between CXCR6 and CD103 as revealed by the highly significant correlation between these two markers. Further, CXCR6 was found associated with increased survival only with CD8 high ovarian cancer patients.

To corroborate the human findings of CXCR6 and resident memory T cells, we used prophylactic mice tumor models. The advantage of using the prophylactic model over the therapeutic model is that it increases the probability of tumor responses to be derived from memory CD8 T cells. This is not possible in the therapeutic model, where the constant presence of tumor mitigates memory formation or makes it dysfunctional. Our results from mice extend the human observations on CXCR6 and Trm cells. Additionally, it suggests that chemokine receptor CXCR6 is not involved in T cell trafficking to tumors, as that is served by CXCR3 which shows high expression on both resident and circulatory memory T cells. This unbiased expression of CXCR3 on both memory subsets is also seen in human samples. In contrast, CXCR6 serves as a retention factor to keep T cells in ovarian peritoneal metastatic sites, increasing their likelihood of becoming resident memory cells. This is supported by the fact that CXCR6 expression is acquired late and occurs *in situ* in the tumor, and CXCR6 is highly expressed in tumor-specific T cells that are resident but not by those in circulation. Strong association of CXCR6 with CD103 (a marker of tissue residency), in both human and mouse studies also attest to the retention role. Further observation of proximity of CD103^+^ T cells to cells that express CXCL16 (the primary chemokine ligand for CXCR6), serves as additional proof for the role of CXCR6 in resident memory T cell localization in ovarian tumor tissues. Characterization of cells that produce CXCL16 and help localize CXCR6^+^ CD103^+^ within the ovarian tumor microenvironment, revealed that the majority of these CXCL16 producing cells were negative for CD45 and EpCAM. This warrants further characterization of these cells in the future using more extensive mass cytometry technologies. Knockout of CXCR6 in tumor-specific T cells enhanced their responses in blood and spleen. However, they showed a reduced response in the tumor as evidenced by low frequencies of CD90.1 TILs, and more specifically, fewer numbers of CD103^+^ resident memory cells in the tumor. This culminated in poor control of tumors by KO T cells in recipient mice. Although we observed some tumor control even in recipient mice of KO T cells, this might be due to some contributory responses from circulatory memory cells, as reported previously (45).

Based on the above results, we hypothesize that DC/tumor antigen-activated T cells express CXCR3, driving T cell infiltration into the tumor or peripheral tissues. Though the current study didn’t verify CXCR3 role in peripheral migration using CXCR3KO studies, previous studies support this fact(46,47). Once in the tumor, *in situ* acquisition of CXCR6 dictates whether T cells will remain localized to the tumor or emigrate from the ovarian peritoneal tissue microenvironment. CXCR6 on tumor-specific T cells then engage with CXCL16 derived from the TME and facilitates T cell retention. The role of CXCR6 in T cell retention is supported by the previous demonstration that the unique DRF motif in CXCR6 makes it more suited for adhesion than chemotaxis to ligand CXCL16 (48) which exists in both transmembrane and soluble forms (49). We envisage that CXCR6 mediated retention may increase the probability of T cell exposure to cytokines like IL-15, TGFβ and other influences in tissue milleu that drive Trm development (36,37,50).

In conclusion, the present study implicates CXCR6 as a critical regulator of residency and persistence of memory CD8 T cell responses in ovarian TME, thereby increasing enhanced immunosurveillance and control of ovarian cancer. This warrants targeting the CXCL16/CXCR6 axis in future immunotherapeutic strategies to improve treatment outcomes for ovarian cancer patients.

## Author Contributions

RM designed and performed the experiments, analyzed results, and wrote the manuscript

AJRM advised, designed, and performed the experiments, reviewed, and corrected the manuscript.

SB and WH did the TCGA and statistical analysis for the current study

AM processed and analyzed samples for the human study.

CE bred/crossed and generated mouse strains used, wrote mice protocols, reviewed data, and manuscript.

BL engineered maraba-OVA viral constructs and revised the manuscript.

PS advised on the experiments, revised the manuscript.

KO designed, advised experiments, reviewed and corrected the manuscript.

## Acknowledgments

We acknowledge the support from grants: NCI SPORE P50CA159981, NIH U24 CA232979, NIH R01 CA158318, NIH U01 CA233085, NCI Cancer Center Support Grant P30 CA016056, NIH/NCI R01CA220297, and NIH/NCI R01CA216426 grants and core facilities: Laboratory animal resource, Flow and confocal core, Translational Imaging Shared Resource for their help with the current study. We acknowledge Dr. Weaver for providing us the T-Lux mice.

## Conflict of interest statement

The authors have declared that no conflicts of interest exist.

## Notes

### Competing Interest Statement

The authors have declared no competing interest.

### Summary of Updates

Title changed The abstract is now structured and updated. Figures consist of only 1-5, 6-8 deleted Accordingly, main text updated Supplemental data updated

