## Supplemental tables and figures for "CXCR6 by increasing retention of memory CD8 T cells in the ovarian tumor microenvironment promotes immunosurveillance and control of ovarian cancer"

**Supplementary Table-1. Primers for SYBR based real-time QPCR analysis of Human genes**

| Gene Name | | Sequence 5'-3' | # CAT | Company |
|---|---|---|---|---|
| CCR1 | Forward | CCTTGGAACCAGAGAGAAG | KSPQ12012 | Sigma-Aldrich |
| | Reverse | AATACCAAGGAGTACAGAGG | " | " |
| CCR2 | Forward | AAGCCTTTTTCACATAGCTC | " | " |
| | Reverse | CTTTCACATTCTTTCCTGGTC | " | " |
| CCR3 | Forward | CACTGCTGAGTTGTATTGG | " | " |
| | Reverse | GCTCTGGTATCAGCTTTTTC | " | " |
| CCR4 | Forward | GCTTTCAGAAAAGCAAGC | " | " |
| | Reverse | TATTCTGTGTAGTGGGATGAG | " | " |
| CCR5 | Forward | TGCTGTTTCTTTTGAAGGAG | " | " |
| | Reverse | TATGCTGGTGAACAGAAATG | " | " |
| CCR6 | Forward | CTCAATAAAGAAGGAGCTGTC | " | " |
| | Reverse | GAACAAAGGCTGTCACTAAG | " | " |
| CCR7 | Forward | AATGATGGAGTACATGATAGGG | " | " |
| | Reverse | CAGACAAGCAAAACAAAGTG | " | " |
| CCR8 | Forward | GAACAAAGGCTGTCACTAAG | " | " |
| | Reverse | GTTTCCCAGAAGACTGAATAC | " | " |
| CCR9 | Forward | GACTAACACAAGCCCTATTC | " | " |
| | Reverse | CACAGTAGAAGTCAGTGAAG | " | " |
| CCR10 | Forward | CTGCGAATCTAGAGGAGG | " | " |
| | Reverse | CACAGAGGTAGTCCCTTTAG | " | " |
| XCR1 | Forward | AGAAACACCAGGCAGTATAG | " | " |
| | Reverse | GACTAACACAAGCCCTATTC | " | " |
| CXCR1 | Forward | TTAAGTCACTCTGATCTCTGAC | " | " |
| | Reverse | TGGTTTGATCTAACTGAAGC | " | " |
| CXCR2 | Forward | CCAGTCAGGATTTAAGTTTACC | " | " |
| | Reverse | GTTGATTTCCAGGGATTCTG | " | " |
| CXCR3 | Forward | GTCCTTGAGGTGAGTGAC | " | " |
| | Reverse | TCTCCATAGTCATAGGAAGAG | " | " |
| CXCR4 | Forward | AACTTCAGTTTGTTGGCTG | " | " |
| | Reverse | GTGTATATACTGATCCCCTCC | " | " |
| CXCR5 | Forward | AGTATCCTCATTTGGGGTAG | " | " |
| | Reverse | GCATTGGATGATTAGGATGG | " | " |
| CXCR6 | Forward | GGTGTTCATCAGAACAGAC | " | " |
| | Reverse | GAAAGACCTTGCTGAACTG | " | " |

| | | | | |
|---|---|---|---|---|
| CXCR7 | Forward | GATGTGGGTTACAAAGCTG | " | " |
| | Reverse | AATCAAATGACCTCCGGG | " | " |
| CX3CR1 | Forward | AAATACCCCATCATTCATGC | " | " |
| | Reverse | TTGTTCCAAACGTTTCTAGG | " | " |
| HPRT1 | Forward | ATAAGCCAGACTTTGTTGG | " | " |
| | Reverse | ATAGGACTCCAGATGTTTCC | " | " |

**Supplementary table 2A. Details of primary and secondary antibodies used for confocal staining of human tumor tissues.**

| **Primary antibodies** | **Target Antigen** | **Antibody clone** | **Isotype** | **Flourophore** | **Cat#** | **Vendor** |
|---|---|---|---|---|---|---|
| | CD103 | EPR4166(2) | Rabbit Monoclonal | None | ab129202 | Abcam |
| | CXCR6 | K041E5 | Mouse IgG2a | None | 356002 | BioLegend |
| | EPCAM | 9C4 | Mouse IgG2b | APC/Fire-750 | 324233 | BioLegend |
| | CD8 | YTC182.20 | Rat IgG2b | None | MCA351G | Bio-Rad |
| **Secondary antibodies** | **Target Antigen** | **Antibody clone** | **Isotype** | **Flourophore** | **Cat#** | **Vendor** |
| | Rabbit IgG | Polyclonal | Goat IgG | Alexa-488 | 4412s | Cell signaling Technology |
| | Rat IgG | Polyclonal | Goat IgG | Alexa-555 | 4417s | Cell Signaling Technology |
| | Mouse IgG | Polyclonal | Goat IgG | Alexa-647 | 4410s | Cell signaling technology |

**Supplementary table 2B. Details of primary and secondary antibodies used for confocal staining of mouse tumor tissues.**

| | **Target Antigen** | **Antibody clone** | **Isotype** | **Fluorophore** | **Cat#** | **Vendor** |
|---|---|---|---|---|---|---|
| **Primary antibodies** | CXCL16 | Polyclonal | Rabbit | None | 119350 | Abcam |
| | CD103 | M290 | Rat IgG2a | A700 | 565529 | BD Biosciences |
| | EP-CAM | G8.8 | Rat IgG2a | Alexa594 | 118222 | BioLegend |
| | CD45 | 30-F11 | Rat IgG2b | Alexa647 | 103124 | BioLegend |
| | F4/80 | BM8 | Rat IgG2a | Alexa594 | 123140 | BioLegend |
| **Secondary antibodies** | **Target Antigen** | **Antibody clone** | **Isotype** | **Flourophore** | **Cat#** | **Vendor** |
| | Rabbit IgG | Polyclonal | Goat IgG | Alexa-488 | 4412s | Cell Signaling Technology |

**A**

| Stage | No of Patients |
|---|---|
| III A | 7 |
| III B | 14 |
| III C | 221 |
| IV | 38 |
| Total | 280 |

**B**

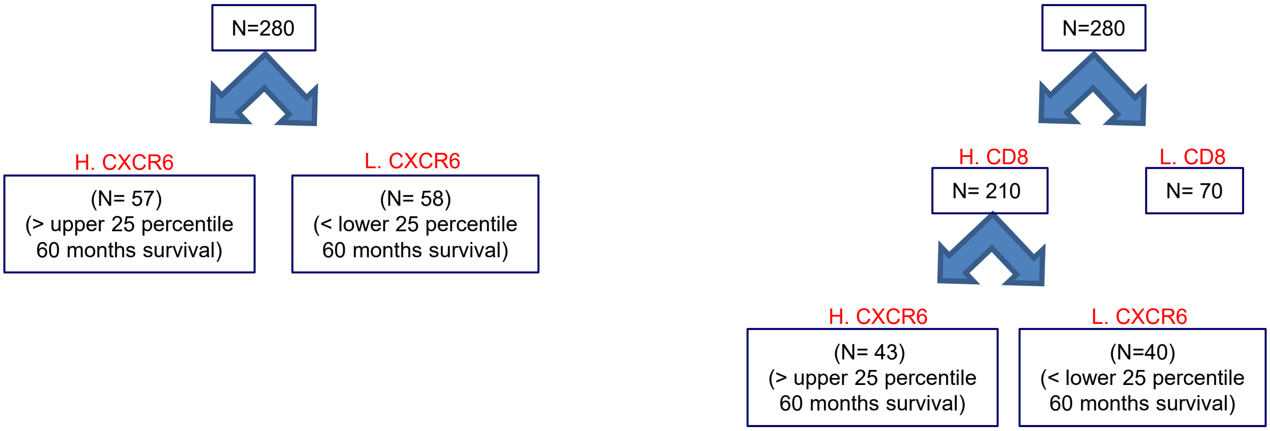


**Supplementary figure 1. Details of high Grade TCGA patient database and stratification. (A)** Info on the stages and no patients in each stage for the TCGA database of Ovarian cancer patients used in the current study. Strategy of sample stratification of TCGA database for to analyze **(B)** correlation and survival with markers CXCR6.

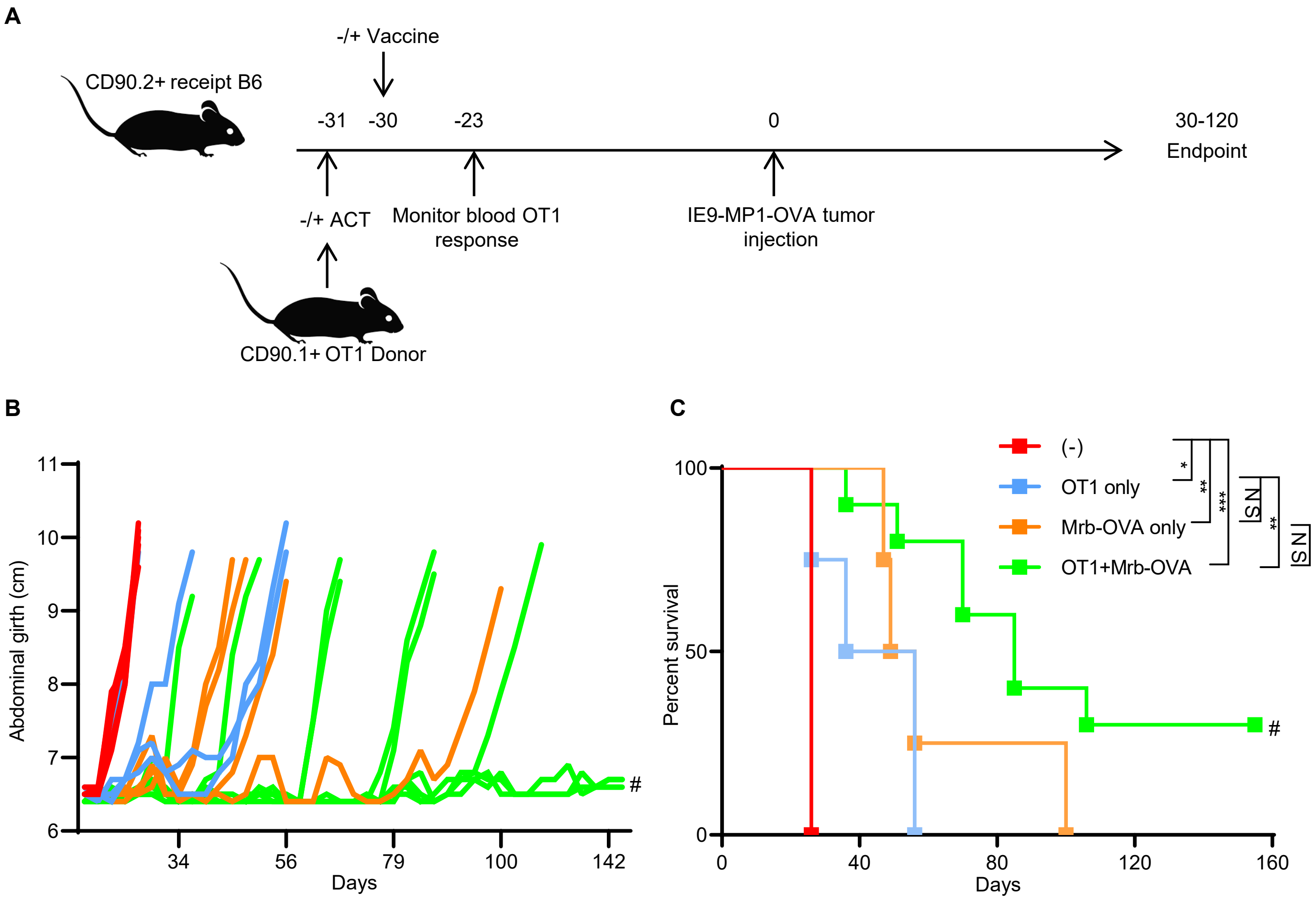


**Supplementary figure 2. Treatment with combination of adoptive transfer OT1 and vaccination with Mrb-OVA offers better protection than single treatments. (A)** Experimental schema of the prophylactic model used to assess test therapeutic efficacy of individual or combination treatments with OT1 T cells and Mrb-OVA vaccine. Tumor progression **(B)** and survival **(C)** of mice treated with individual or combination of OT1 T cells and Mrb-OVA vaccine.

**A**

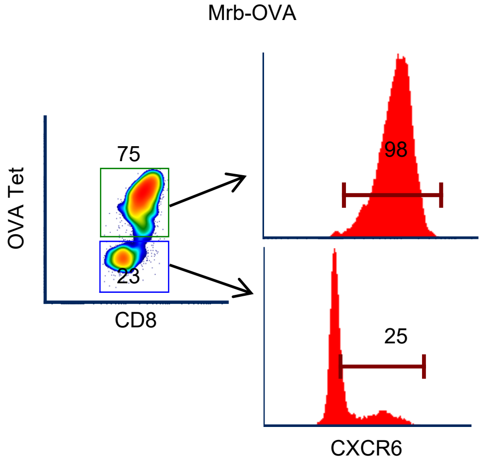


**B**

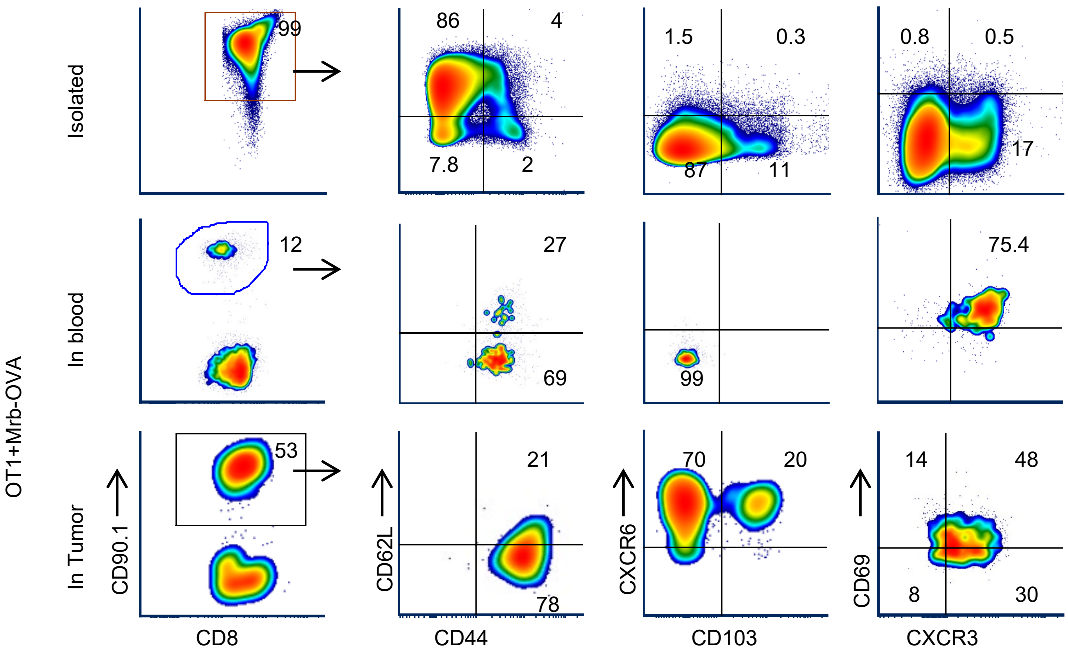


**Supplementary figure 3. CXCR6 marks tumor specific resident memory cells**. **(A)** CXCR6 expression in tumor specific (OVA tetramer pos) and non-tumor (OVA tetramer neg) CD8+ T cells in tumors of mice treated with Maraba-OVA vaccination (Mrb-OVA). **(B)** phenotype of OT1 T cells that were freshly isolated (top panel) from the donor mice or 7 days after vaccination in blood (middle panel) and in endpoint tumors (bottom panel) of mice treated with adoptive cell transfer of OT1 T cells and Maraba-OVA vaccination (OT1+Mrb-OVA).

**A**

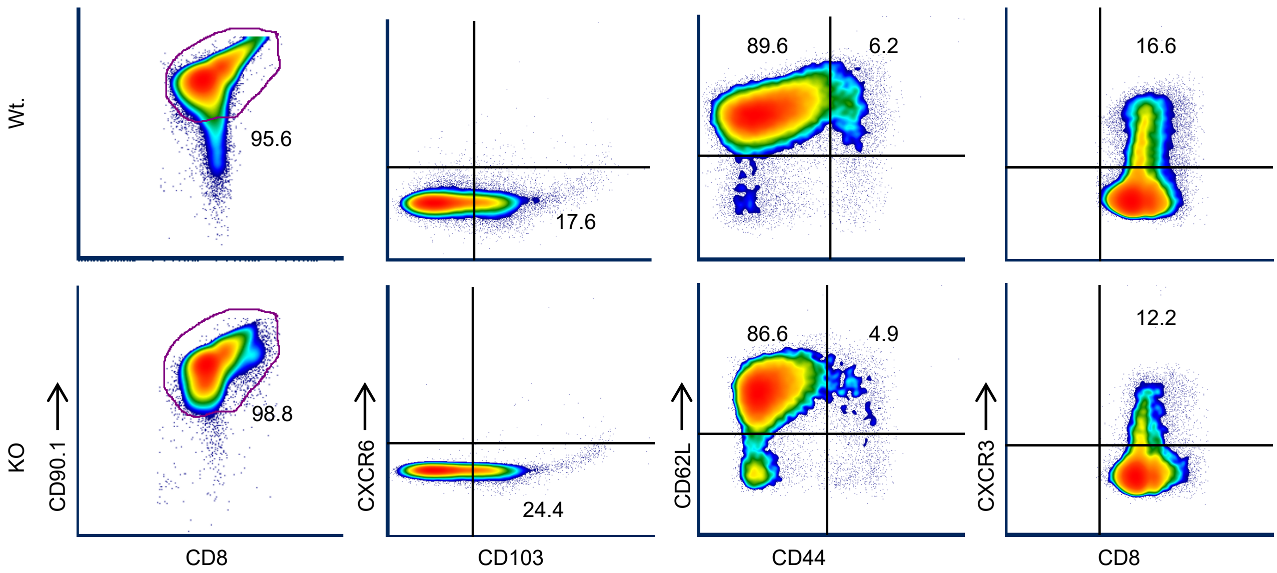


**B**

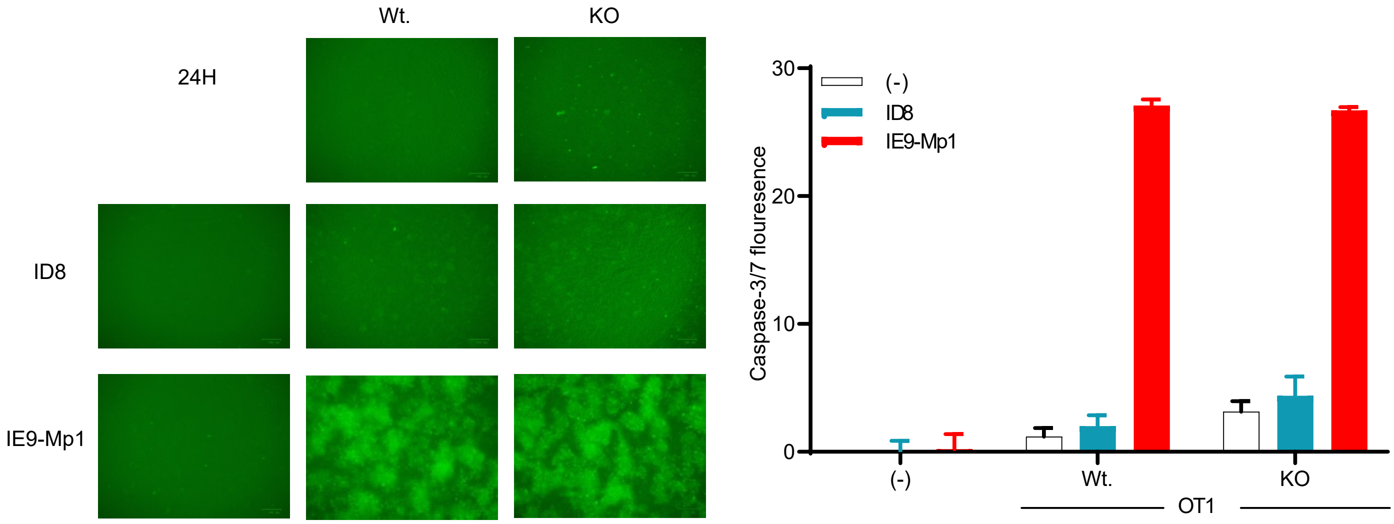


**Supplementary figure 4. CXCR6 knockout did not affect phenotype or killing function of OT1 T cells. (A)** Phenotype analysis of wild (Wt.) or CXCR6KO (KO) OT1 that are freshly isolated from mice for transfer into recipient mice. **(B)** Ability to Wt. or KO OT1 cells to induce activated CASPASE-3/7 in OVA expressing IE9-Mp1 or non-expressing ID8 control in 24-hour co-culture at effector: target ratio 10:1. Left panel shows fluorescence images taken on and right bar graphs shows mean of 3 replicate fluorescence values of various single or co cultures as measured on Biotek Synergy HT microplate plate reader at emission 528nM

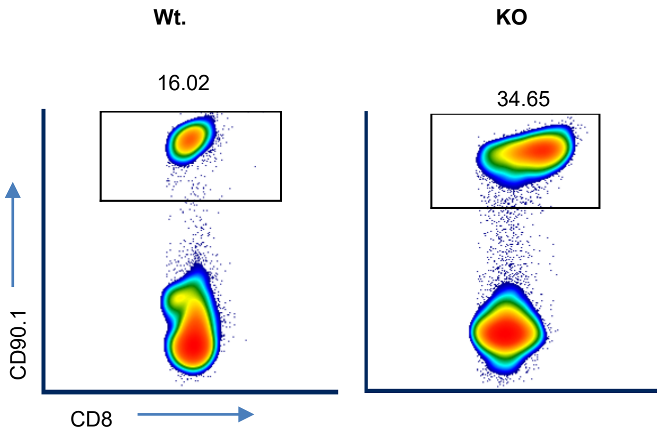


**Supplementary figure 5. CXCR6KO OT1 accumulate more than wt. OT1 cells in spleens of B6 recipient mice.** FACS dot blots of % CD90.1$^{+}$ T cells in spleen of **o**ne representative B6 recipient mice that received ACT of either Wt. or KO OT1 cells +Mrb-OVA vaccine.
